# RNA promotes MAPK signaling through direct interaction with BRAF

**DOI:** 10.64898/2026.10.22.760349

**Authors:** Alexia Le Barch, Sabrina Mennour, Lucilla Fabbri, Titash Sen, Isabelle Girault, Virginie Quidville, Jessica Andreani, Raphael Guérois, Caroline Robert, Patricia Uguen, Stéphan Vagner

## Abstract

RNAs are increasingly recognized as direct regulators of protein function, yet whether RNA can control the activation of signaling kinases remains poorly understood. Here we identify RNA as a direct regulator of BRAF, a central kinase in the RAS-RAF-MEK-ERK (MAPK) pathway. Using complementary crosslinking and RNA-protein interaction approaches, we show that BRAF directly binds RNA in cells and that this interaction is enhanced upon MAPK pathway activation. Depletion of intracellular RNA reduces BRAF:CRAF complex assembly and impairs MEK1/2 phosphorylation, indicating that RNA contributes to RAF complex formation and MAPK signal transduction. We identify a positively charged surface within the BRAF kinase domain that mediates RNA binding. A separation-of-function RNA-binding mutant selectively attenuates BRAF-dependent MEK1/2 phosphorylation, demonstrating that RNA binding is required for efficient MAPK signaling. Moreover, artificial recruitment of the RNA-binding-deficient BRAF mutant to an RNA scaffold restores its signaling activity, supporting a direct role for RNA-mediated BRAF complex assembly. These data identify RNA as an active component of RAF kinase regulation and establish RNA-protein interactions as an additional determinant of MAPK pathway activation.

## INTRODUCTION

RNAs are best known for their roles in gene expression, but accumulating evidence indicates that they can also directly regulate protein function, extending their influence beyond canonical RNA-mediated processes (Hentze et al. 2025). RNA binding can alter protein conformation, catalytic activity, substrate specificity and post-translational modification, as well as regulate protein–protein interactions and the assembly of macromolecular complexes (Abe et al. 2023; Du et al. 2016; Lee et al. 2016; Markaki et al. 2021; Zhu et al. 2022). Thus, RNA can function not only as a template or regulatory intermediate, but also as a dynamic molecular organizer that directly controls protein activity and macromolecular assembly. Whether such mechanisms contribute to the regulation of cellular signaling pathways remains largely unexplored.

The RAS–RAF–MEK–ERK (MAPK) pathway is a central regulator of cell proliferation, differentiation, metabolism and survival and is frequently dysregulated in cancer (Sanchez-Vega et al. 2018; Sinkala et al. 2021). A key step in pathway activation is the regulated dimerization of RAF kinases following RAS activation, which promotes RAF kinase activity and subsequent phosphorylation of MEK1/2. Although the molecular mechanisms governing RAF activation have been extensively studied, whether RNA contributes directly to this process is unknown.

Here we show that BRAF directly binds RNA and that RNA binding promotes BRAF signaling. Our data therefore reveals an additional layer of control over MAPK pathway activation involving an RNA-protein interaction.

## RESULTS

### BRAF interacts with RNA upon MAPK pathway activation

To examine direct RNA binding by BRAF, we used crosslinking immunoprecipitation (CLIP) experiments (Ule et al. 2005, 2003). Briefly, living cells were UV-C crosslinked to covalently stabilize RNA–protein interactions before cell lysis and partial RNA digestion. Following immunoprecipitation, protein-bound RNA fragments were radiolabeled with T4 polynucleotide kinase and [γ-^32^P]-ATP and resolved by denaturing SDS–PAGE. A radioactive signal at the molecular weight of the immunoprecipitated protein indicates direct RNA binding.

CLIP assays were performed in R-Malme-3M melanoma cells which harbor a heterozygous BRAF (wild-type/V600E) genotype (Boussemart et al. 2014). This genotype recapitulates the genetic configuration of most BRAF-mutant melanomas, in which oncogenic BRAF V600E is co-expressed with wild-type (WT) BRAF. The CLIP experiment revealed a radioactive signal migrating at the molecular weight of BRAF in samples irradiated with UV and treated with RNase A (hereafter referred to as RNase), a single-stranded RNA-specific endoribonuclease that degrades RNA by cleaving after cytidine and uridine residues (Fig. 1A). No signal was detected in the absence of UV crosslinking, indicating that the signal reflects bona fide RNA–protein crosslinks rather than nonspecific RNA association (Fig. 1A). As expected, increasing RNase concentration progressively sharpened and intensified the signal, consistent with trimming of the crosslinked RNAs to short fragments that remain covalently attached to the protein (Fig. 1A). Efficient BRAF immunoprecipitation in both UV-crosslinked and non-crosslinked samples excluded differential protein recovery as the basis for the UV-dependent signal (Fig. 1A). Moreover, siRNA-mediated depletion of BRAF markedly reduced the radioactive signal, confirming its identity as a BRAF-RNA complex (Fig. 1B). Together, these data indicate that BRAF, comprising WT and/or V600E species, directly associates with RNA.

**Figure 1.**
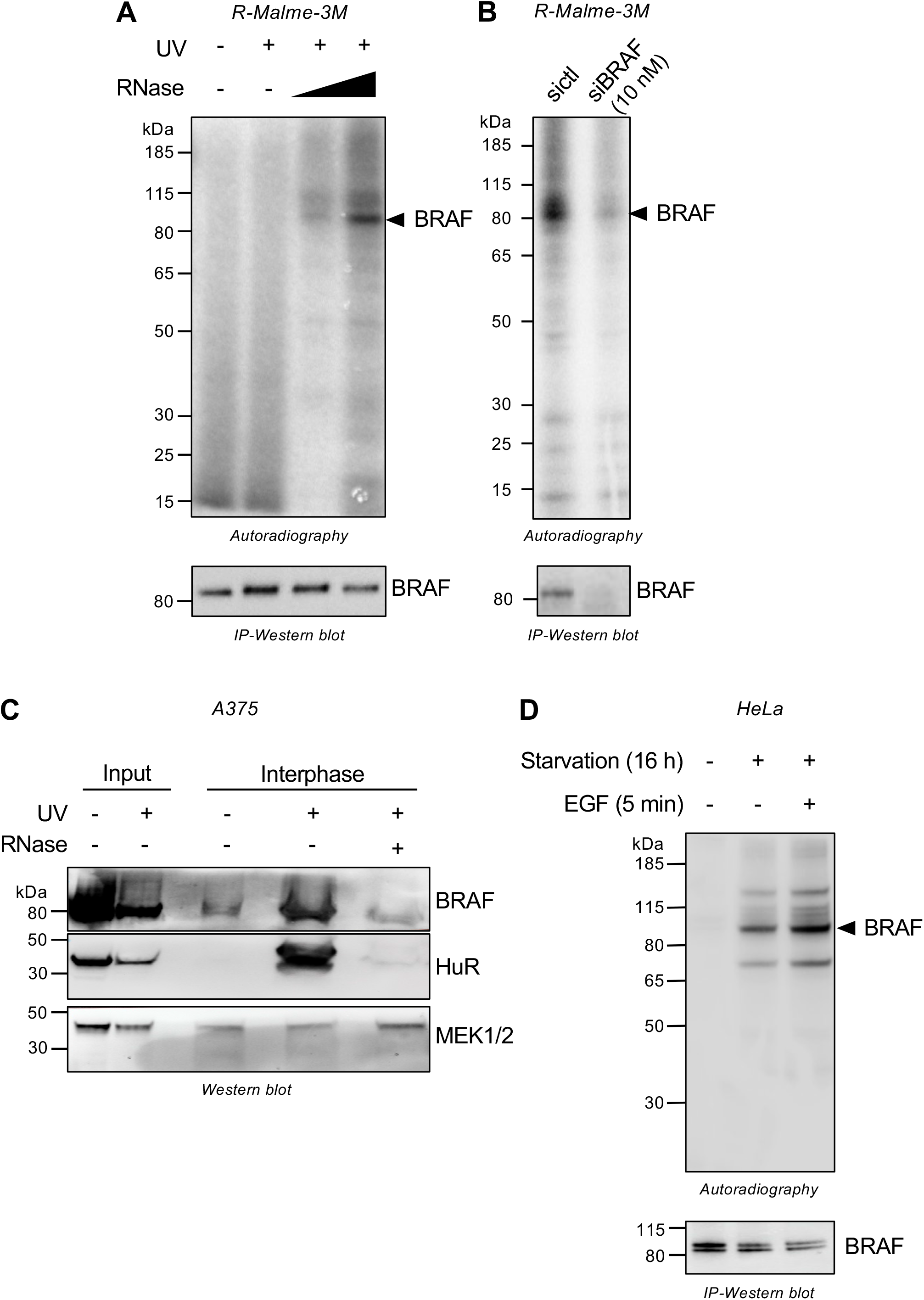
BRAF interacts with RNA upon MAPK pathway activation. **(A**) Crosslinking immunoprecipitation (CLIP) of endogenous BRAF in R-Malme-3M cells. Upper panel: Autoradiography of BRAF–RNA complexes from UV-C (UV)-irradiated (+) or non-irradiated (−) cells. Two RNase concentrations were applied prior to immunoprecipitation performed in stringent conditions. A prominent band corresponding to the expected molecular weight of BRAF is indicated by a black arrow. Lower panel: Immunoblot of BRAF immunoprecipitates under the same experimental conditions shown in the upper panel. **(B)** CLIP of endogenous BRAF in R-Malme-3M cells transfected with an siRNA targeting BRAF (siBRAF) or a control siRNA (sictl). Upper panel: Autoradiography of BRAF-RNA complexes from UV-irradiated cells. A prominent band corresponding to the expected molecular weight of BRAF is indicated by a black arrow. Lower panel: Immunoblot of BRAF immunoprecipitates under the same experimental conditions. **(C)** Immunoblot analysis of orthogonal organic phase separation (OOPS) fractions from UV-irradiated (+) or non-irradiated (−) A375 cells. RNase A treatment (+) of cell lysate prior to loading to the column is used as a negative control. The RBP HuR was used as a positive control and MEK1/2 as a negative control. **(D)** CLIP of endogenous BRAF in HeLa cells. Upper panel: Autoradiography of BRAF-RNA complexes from UV-irradiated cells, serum-starved (+) or not (-) for 16 h and stimulated (+) or not (-) with EGF for 5 min. A prominent band corresponding to the expected molecular weight of BRAF is indicated by a black arrow. Lower panel: Immunoblot of BRAF immunoprecipitates under the same experimental conditions.

To independently validate the CLIP results, we used orthogonal organic phase separation (OOPS), an antibody-independent method that isolates UV-crosslinked RNA–protein complexes by phenol–chloroform partitioning (Queiroz et al. 2019). In A375 melanoma cells, which are homozygous for the V600E mutation, BRAF V600E was enriched in the RNA-bound fraction (interphase) following UV crosslinking, with an enrichment comparable to that of the canonical RNA-binding protein HuR (Fig. 1C). This enrichment was abolished by RNase treatment prior to OOPS, whereas MEK1/2 was not recovered in the RNA-bound fraction under any condition (Fig. 1C). Together, these orthogonal approaches establish that oncogenic BRAF V600E directly binds RNA.

Although BRAF has not emerged as a candidate RBP in initial RNA interactome studies (Baltz et al. 2012; Beckmann et al. 2016; Castello et al. 2012), those analyses were not performed in cancer cells in which the MAPK pathway is active. One study, however, defined the RNA binding proteome of a NRAS-mutated melanoma cell line, but did not identify BRAF (Mestre-Farràs et al. 2022). We nevertheless asked whether RNA binding by BRAF is induced by MAPK pathway activation. Cells were serum-starved for 16 h to minimize basal receptor tyrosine kinase activity and enable synchronous activation of the MAPK pathway upon EGF stimulation. A 5 min-long EGF stimulation was selected for subsequent analyses, as it led to robust MEK1/2 phosphorylation (Fig. S1). CLIP analysis of serum-starved HeLa cells revealed a BRAF-associated radioactive signal that increased following EGF stimulation (Fig. 1D). Comparable BRAF recovery in both conditions excluded differences in immunoprecipitation efficiency (Fig. 1D). These data indicate that MAPK pathway activation enhances RNA binding by WT BRAF.

To determine whether MAPK pathway activation broadly regulates RNA binding by BRAF, we performed CLIP in Flp-In-293 cells stably expressing V5-tagged BRAF. Expression of oncogenic NRAS Q61K, which induced MEK1/2 phosphorylation, markedly increased the UV-dependent radioactive species corresponding to BRAF relative to expression of NRAS that failed to activate the pathway (Fig. S2A). To exclude the possibility that the ^32^P signal reflected protein phosphorylation rather than RNA crosslinking, the radioactive species were excised from the gel, subjected to proteinase K digestion, and the released material was resolved by denaturing TBE-Urea gel electrophoresis. Radiolabeled ∼100-nt RNA fragments were recovered after protein digestion and were eliminated by RNase treatment, confirming that the detected signal originated from RNA covalently crosslinked to BRAF rather than from protein phosphorylation (Fig. S2B). Together, these data demonstrate that direct RNA binding by BRAF is dynamically enhanced upon MAPK pathway activation.

### Intracellular RNA promotes BRAF:CRAF complex assembly and MAPK pathway activation

Given that RNA can promote the assembly and/or stabilization of protein complexes, we next explored the possibility that the interaction of BRAF with RNA contributes to RAF complex regulation during MAPK pathway activation (Fig. 2A). To investigate whether intracellular RNA influences BRAF:CRAF association, we monitored BRAF:CRAF proximity using a proximity ligation assay (PLA) following RNase-mediated RNA depletion in permeabilized HeLa cells. EGF stimulation increased BRAF:CRAF PLA signals, consistent with activation-dependent assembly of RAF complexes (Fig. 2B-C). RNase treatment of permeabilized cells markedly reduced BRAF:CRAF proximity (Fig. 2B-C) without affecting BRAF or CRAF protein abundance (Fig. S3A-B), indicating that intracellular RNA contributes to the formation or stabilization of RAF complexes.

**Figure 2.**
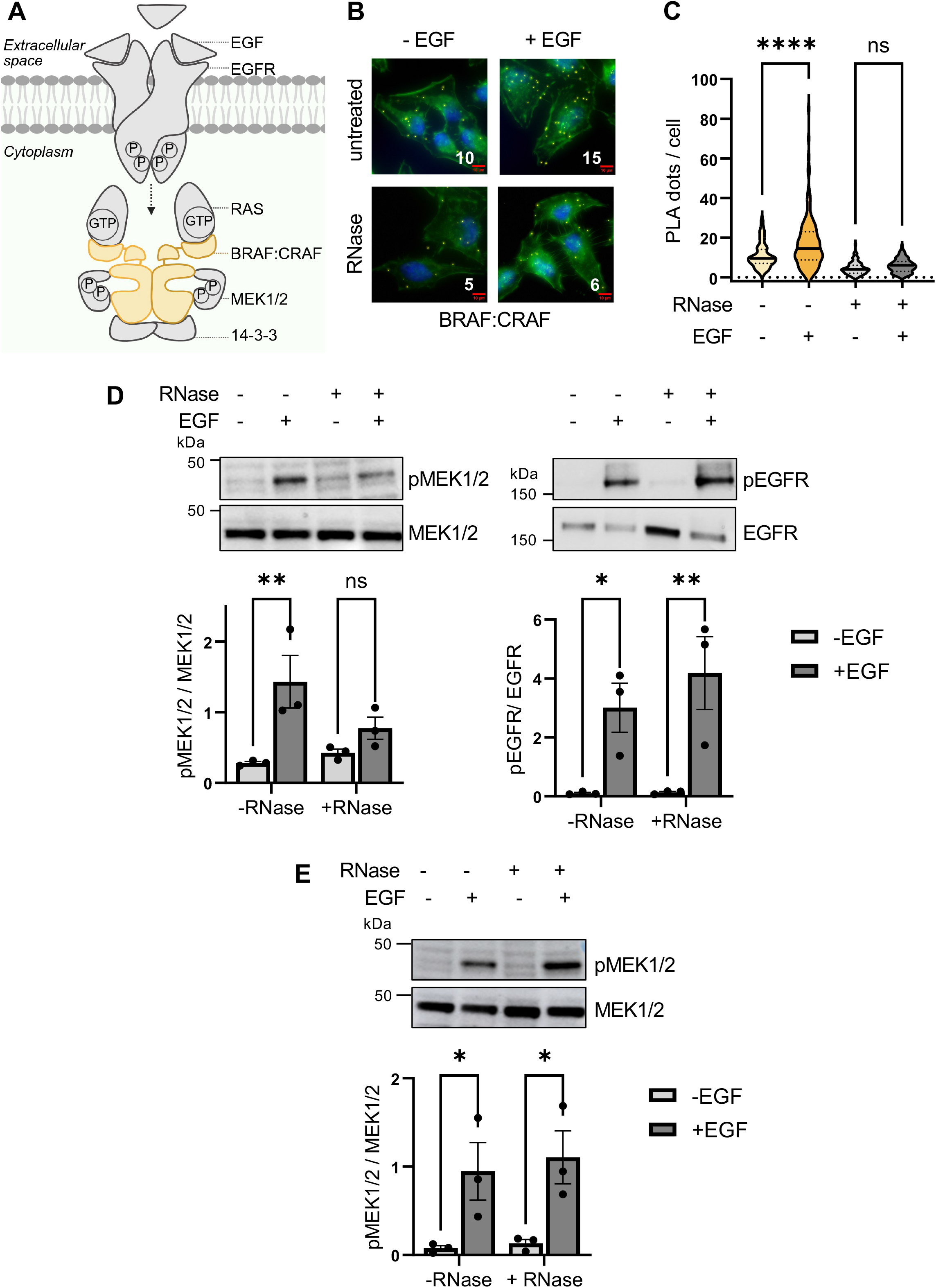
Intracellular RNA promotes BRAF:CRAF complex assembly and enhances MAPK pathway activation. **(A)** Schematic representation of BRAF dimer-mediated MAPK signal transduction. Binding of EGF induces EGFR dimerization and autophosphorylation, leading to the recruitment of RAS to the plasma membrane, where it becomes activated through GTP binding. Active RAS promotes the recruitment and dimerization of BRAF and CRAF. The BRAF heterodimer is stabilized by two 14-3-3 chaperone proteins and subsequently phosphorylates MEK1/2, propagating MAPK signaling. The dashed arrow represents the intermediate adaptor proteins necessary for RAS activation, which are omitted here to simplify the schema. **(B)** Representative images of proximity ligation assays (PLA) measuring BRAF:CRAF proximity in HeLa cells stimulated with EGF for 5 min (+) or left unstimulated (-). Cells were permeabilized, treated with (+) or without (-) RNase A prior to fixation, and processed for PLA. BRAF:CRAF PLA signal (yellow), phalloidin (green), and DAPI (blue). Scale bar, 10 μm (red). **(C)** Quantification of three independent PLA experiments shown as violin plots (> 100 cells in total for each condition). Untreated samples are shown in shades of yellow, and RNase-treated samples in shades of gray. Statistical significance was assessed by ordinary one-way ANOVA followed by Tukey’s multiple-comparisons test. Of the six pairwise comparisons, only the comparisons between unstimulated and EGF-stimulated cells within each treatment group are indicated (−RNase: −EGF vs +EGF, \*\*\*\**P* < 0.0001; +RNase: −EGF vs +EGF, *P* = 0.5906). **(D)** Immunoblot analysis of serum-starved (16 h) HeLa cells stimulated with EGF for 5 min (+) or left unstimulated (-), permeabilized and subsequently treated with (+) or without (-) RNase A. Histograms showing the quantification of phosphorylated-to-total protein ratios from three independent experiments are shown below each immunoblot. P values were determined using 2-way ANOVA followed by Sidak’s multiple-comparisons for pMEK1/2 (−RNase: −EGF vs +EGF, \*\**P* = 0.0079; +RNase: −EGF vs +EGF, *P* = 0.4535) and pEGFR (−RNase: −EGF vs +EGF, \**P* = 0.0489; +RNase: −EGF vs +EGF, \*\**P*= 0.0098). **(E)** Immunoblot analysis of non-permeabilized, serum-starved (16 h) HeLa cells stimulated with EGF for 5 min (+) or left unstimulated (-), following treatment with (+) or without (-) RNase A (RNase), as described in (D). P values were determined using 2-way ANOVA followed by Sidak’s multiple-comparisons for pMEK1/2 (−RNase: −EGF vs +EGF, \**P* = 0.0485; +RNase: −EGF vs +EGF, \**P* = 0.0295).

We next examined the effect of RNA depletion on MEK1/2 phosphorylation. RNase treatment of permeabilized cells specifically impaired MEK1/2 phosphorylation without affecting EGFR phosphorylation (Fig. 2D). Since extracellular RNAs have been implicated in intercellular signaling (Chai et al. 2026), we next asked whether they contribute to MAPK pathway activation. RNase treatment of non-permeabilized cells, thereby restricting RNA degradation to the extracellular compartment, had no detectable effect on EGF-induced MEK1/2 phosphorylation (Fig. 2E). These results exclude a measurable contribution of extracellular RNAs under these experimental conditions. Together, these data indicate that intracellular RNAs contribute to the regulation of MAPK signaling downstream of EGFR activation and upstream of MEK1/2.

### An RNA-binding-deficient BRAF mutant impairs MAPK signal transduction

Although RNase-mediated RNA depletion demonstrated that intracellular RNA regulates RAF complex assembly and MEK1/2 phosphorylation (Fig. 2), this approach does not distinguish between a direct role for RNA binding by BRAF and indirect effects arising from the disruption or recruitment of other RNA-protein interactions. To directly assess the functional significance of the BRAF-RNA interaction, we sought to generate an RNA-binding-deficient BRAF mutant.

Inspection of the highest-resolution BRAF structure comprising dimers of both BRAF and 14-3-3 (Liau et al. 2020) identified a surface-exposed patch in the C-terminal region of the kinase domain enriched in conserved basic residues (R719, K723, and R726) (Fig. 3A), suggesting a candidate RNA-binding interface. We therefore generated charge-altering mutants in which these residues were substituted in combination with either alanine (3A), or glutamate (3E) or individually (A1, A2, A3) (Fig. 3B). The RNA-binding capacity of these variants was assessed by CLIP following transient expression of GFP-tagged BRAF proteins in HEK293T cells.

**Figure 3.**
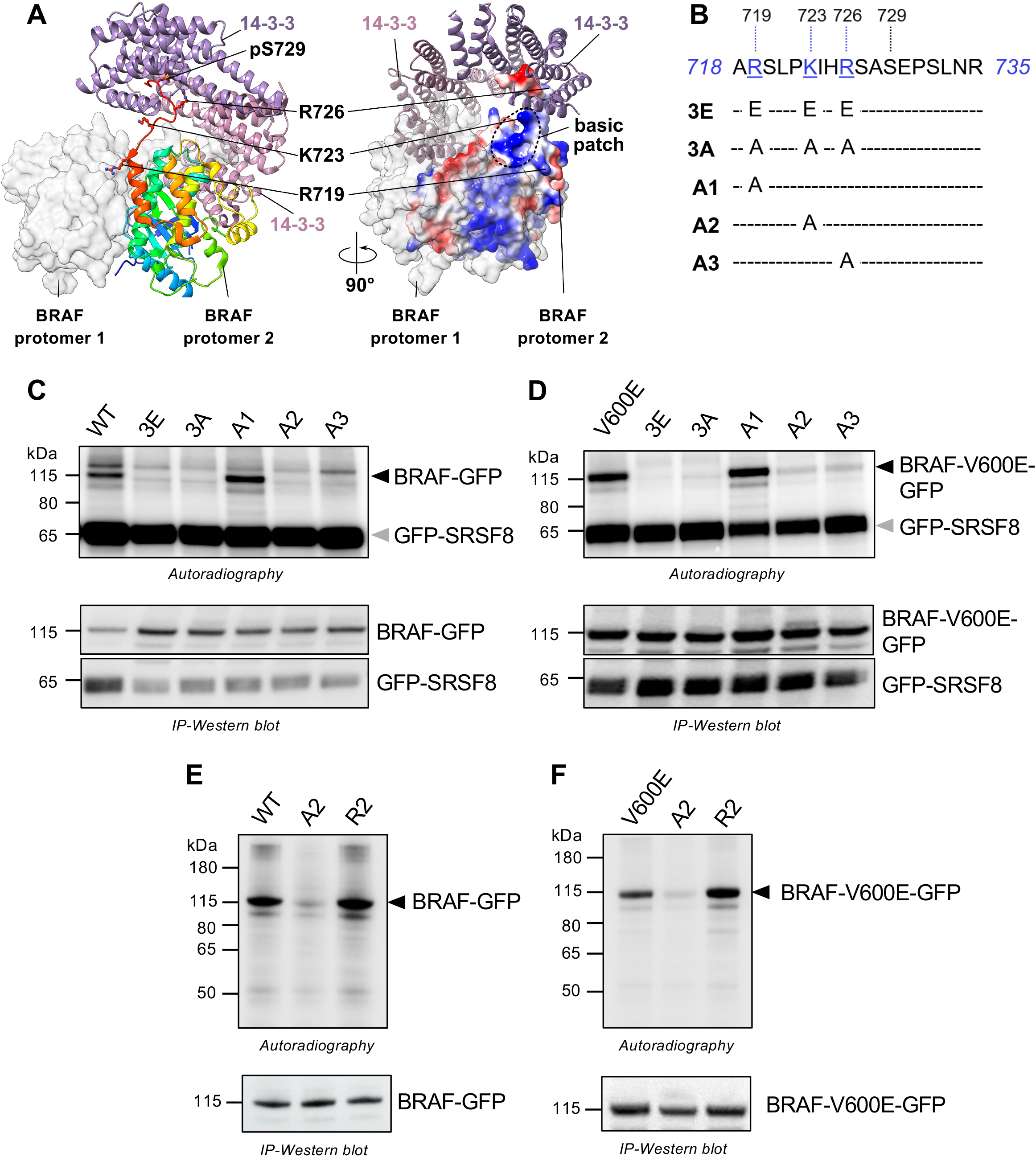
Identification of RNA-binding-deficient BRAF mutants within a positively charged surface region. **(A)** Structural and electrostatic properties of BRAF dimer bound to a tandem of 14-3-3 domains based on the crystal structure of the apo BRAF dimer bound to a 14-3-3 dimer (PDB 6XAG). Left panel, BRAF protomer 1 is represented as a white molecular surface, whereas BRAF protomer 2 is shown as a cartoon colored from the N- to the C-terminus (blue to red). The two 14-3-3 protomers are shown in purple and pink. The C-terminal region of BRAF protomer 2 containing residues R719, K723, R726, and phosphorylated S729 (pS729) is highlighted with pS729 engaged in the phosphoserine-binding groove of 14-3-3. Right panel, a 90°-rotated view shows the electrostatic surface potential of BRAF protomer 2 with positively charged regions in blue and negatively charged regions in red. The positively charged surface region adjacent to the C-terminal BRAF is indicated as the basic patch. Molecular graphics and structural analyses were performed using UCSF ChimeraX (Pettersen et al. 2021). **(B)** Amino acid sequence of the C-terminal region of BRAF highlighting the positively charged residues (blue) and the studied mutations targeting them. **(C)** CLIP of BRAF-GFP with the indicated mutations and GFP-SRSF8 (used as a positive control for crosslinking efficiency and GFP immunoprecipitation recovery) from co-transfected HEK293T cells. Upper panel: Autoradiography of BRAF-RNA and SRSF8-RNA complexes from UV-irradiated cells. Prominent bands corresponding to the expected molecular weight of the BRAF-GFP-RNA and GFP-SRSF8-RNA complexes are indicated with a black and a grey arrow, respectively. Lower panel: Immunoblot of GFP immunoprecipitates under the same experimental conditions. **(D)** CLIP experiments, as described in (C), with BRAF V600E-GFP instead of BRAF-GFP. **(E)** CLIP of BRAF-GFP with the indicated mutations from transiently transfected HEK293T cells. Upper panel: Autoradiography of BRAF-RNA complexes from UV-irradiated cells. A prominent band corresponding to the expected molecular weight of BRAF-RNA complexes is indicated with a black arrow. Lower panel: Immunoblot of GFP immunoprecipitates under the same experimental conditions. **(F)** CLIP experiments, as described in (E), with BRAF V600E-GFP instead of BRAF-GFP.

Compared with BRAF, the 3E, 3A, A2, and A3 mutants displayed markedly reduced RNA binding, whereas the A1 mutant retained activity (Fig. 3C). GFP–BRAF immunoprecipitation efficiency was comparable across all variants, excluding altered protein recovery as the basis for reduced CLIP signals. Moreover, co-expression of GFP-tagged SRSF8, an established RNA-binding protein, yielded comparable CLIP signals across conditions, confirming consistent crosslinking and recovery efficiency (Fig. 3C). Introducing the same mutations into oncogenic BRAF V600E produced an equivalent pattern of RNA-binding defects (Fig. 3D). Furthermore, replacing the critical lysine residue (K723) with arginine, thereby preserving positive charge, did not affect RNA binding in both WT and BRAF V600E backgrounds (Fig. 3E-F). Together, these results identify a positively charged surface on BRAF that directly contributes to RNA binding.

We next investigated the functional consequences of mutating the basic patch on BRAF signaling. HEK293T cells were transiently expressing BRAF or RNA-binding-deficient BRAF mutants, and MEK1/2 phosphorylation was assessed (Fig. 4A). Mutations within the basic patch that affect RNA binding (3E, 3A, A2, and A3), but not the one that retained RNA binding (A1) reduced MEK1/2 phosphorylation relative to total MEK1/2 protein levels, suggesting that RNA binding is involved in BRAF signaling.

**Figure 4.**
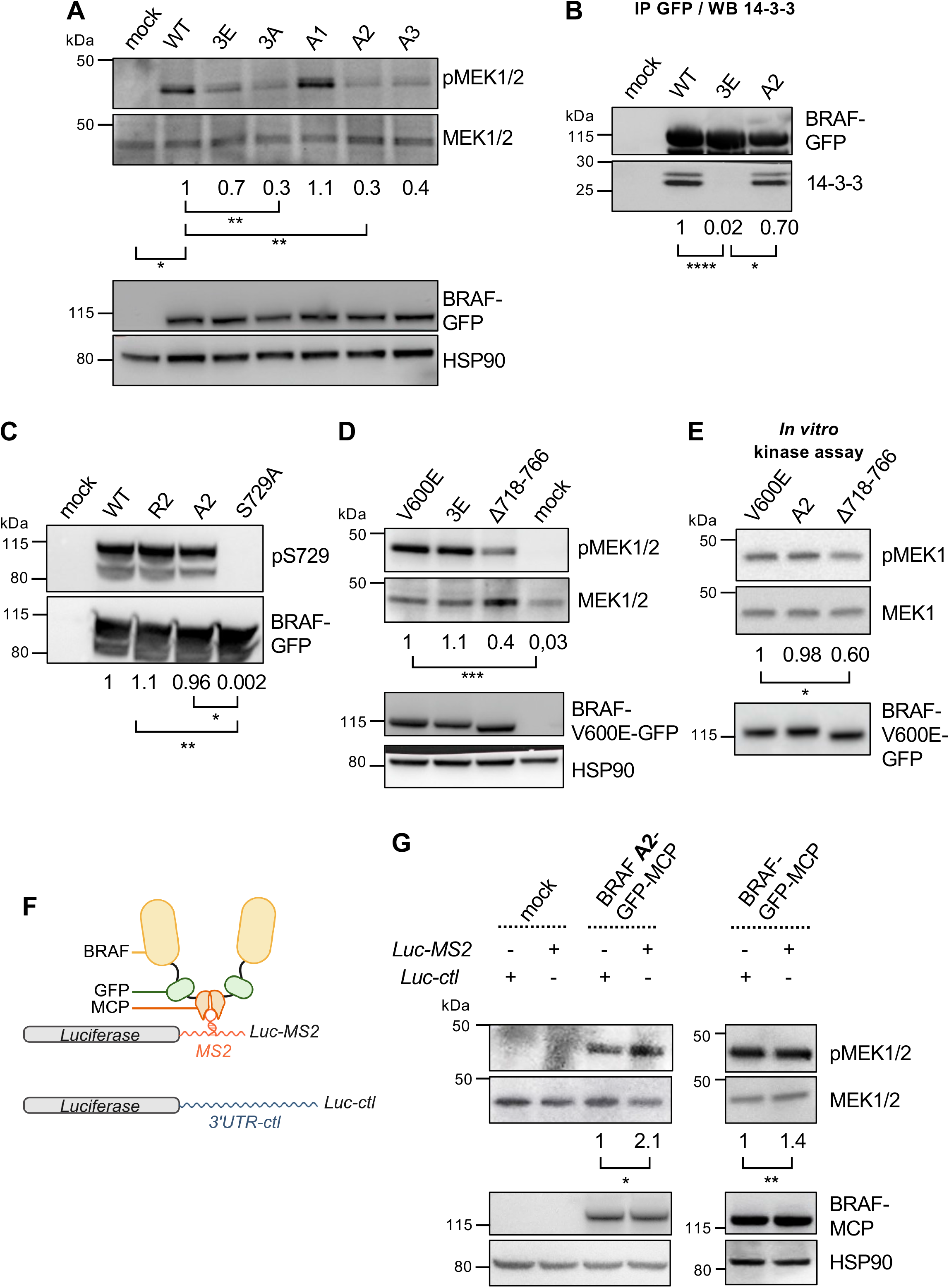
An RNA-binding-deficient BRAF mutant impairs MAPK signaling. **(A)** Immunoblot analysis of BRAF-GFP with the indicated mutations from transiently transfected HEK293T cells following FACS-based sorting for equivalent GFP expression. Phosphorylated-to-total protein ratios were quantified from three independent experiments, and the fold changes for the indicated BRAF mutants were calculated relative to BRAF. P values were determined using RM one-way ANOVA followed by Dunnett’s multiple-comparisons (WT vs mock, *\*P* = 0.0196; WT vs 3E, *P* =0.1151; WT vs 3A, *\*\*P* = 0.0052; WT vs A1, P = 0.5828; WT vs A2, *\*\*P* = 0.0094; WT vs A3, P = 0.0664). The median phosphorylated-to-total protein ratio fold change relative to BRAF is indicated below each immunoblot. HSP90 was used as a loading control. **(B)** Immunoblot analysis of Co-IP experiments performed on transiently transfected HEK293T. BRAF-GFP with the indicated mutations was immunoprecipitated and 14-3-3 presence was assessed in immunoprecipitated samples. 14-3-3 - to-BRAF-GFP ratios were quantified from three independent experiments, and the fold changes for the indicated BRAF mutants were calculated relative to BRAF. P values were determined using RM one-way ANOVA followed by Tukey’s multiple-comparisons (WT vs 3E, *\*\*\*\*P* < 0.0001; A2 vs 3E, *\*P* = 0.0363; WT vs A2, *P* = 0.1177). The median 14-3-3-to-BRAF-GFP ratio fold change relative to WT control condition is indicated below each immunoblot. **(C)** Immunoblot analysis, as described in (A). Phosphorylated-to-total protein ratios were calculated from three independent experiments, and the fold changes for the indicated BRAF mutants were calculated relative to BRAF. P values were determined using ordinary one-way ANOVA followed by Tukey’s multiple-comparisons (R2 vs S729A, *\*\*P* = 0.0054; A2 vs S729A, *\*P* = 0.0110; R2 vs A2, *P* = 0.7694). The median phosphorylated-to-total protein ratio fold change relative to WT control condition is indicated below each immunoblot. **(D)** Immunoblot analysis of BRAF V600E-GFP and indicated mutants, as described in (A). P values were determined using RM one-way ANOVA followed by Dunnett’s multiple-comparisons (V600E vs V600E 3E, *P* = 0.9861; V600E vs V600E Δ718-766, *P*=0.2474; V600E vs mock, \**\*\*P* = 0.0009). **(E)** Immunoblot analysis of pMEK1 in *in vitro* kinase assay of BRAF V600E-GFP and indicated mutants immunoprecipitated from transiently transfected HEK293T. pMEK1 level was quantified using RM one-way ANOVA followed by Dunnett’s multiple-comparisons (V600E vs V600E A2, *P* = 0.8552; V600E vs V600E Δ718-766, *\*P* = 0.0330). The median pMEK1 ratio fold change relative to BRAF V600E is indicated below each immunoblot. **(F)** Schematic representation of the synthetic RNA recruitment strategy. Fusion of BRAF–GFP to the bacteriophage MS2 coat protein (MCP) enables dimeric recruitment of BRAF to one cognate MS2 stem-loop sequences inserted into the 3′ UTR of a luciferase transcript (Luc-MS2). A luciferase transcript containing an unrelated 212-nt sequence in place of the MS2 stem loop (Luc-ctl) was used as a negative control. **(G)** Immunoblot analysis from HEK293T cells transiently co-transfected with BRAF A2-GFP-MCP or BRAF-GFP-MCP and either Luc-MS2 or Luc-ctl as indicated. Cells were FACS-based sorted for equivalent GFP expression. Phosphorylated-to-total protein ratios were quantified from three independent experiments, and the fold changes for the Luc-MS2 conditions were calculated relative to Luc-ctl conditions. P values were determined using paired two-tailed Student’s *t*-tests for pMEK1/2 (A2: Luc-MS2 vs Luc-ctl, \**P* = 0.0106; WT: Luc-MS2 vs Luc-ctl, \*\**P* = 0.0028). The median phosphorylated-to-total protein ratio fold change for Luc-MS2 relative to Luc-ctl is indicated below each immunoblot. Mock: untransfected cells. HSP90 was used as a loading control

Although these data are consistent with a role for RNA binding in promoting BRAF signaling activity, we cannot exclude that this conserved basic surface contributes to BRAF function through additional structural or protein–protein interactions. We therefore sought to identify a mutant that preserves the catalytic and protein-interacting properties of BRAF while selectively disrupting its ability to bind RNA. Such a mutant would enable us to directly determine whether RNA binding by BRAF is required for efficient RAF complex assembly and MAPK signal transduction. We focused on the A2 mutant (K723A), as K723 is highly solvent-exposed and displays substantial local mobility, while making no direct contacts with 14-3-3 or with surrounding structural elements (Fig S4A) (Liau et al. 2020). This makes K723 particularly suitable for testing RNA-binding function while minimizing potential perturbations of the 14-3-3 interaction and 14-3-3-dependent RAF regulation. (Fig S4B) (Park et al. 2019) and therefore provides an opportunity to uncouple RNA-binding defects from potential effects on 14-3-3-dependent RAF regulation. Of note, sequence alignment also revealed that this lysine is conserved in CRAF and ARAF (Fig. S4C).

To exclude the possibility that the A2 mutation affects BRAF function by altering 14-3-3 association, we assessed 14-3-3 binding by co-immunoprecipitation and monitored phosphorylation of BRAF at S729, a residue required for C-terminal 14-3-3 binding. Although 14-3-3 association with BRAF 3E was abolished, BRAF A2 retained substantial 14-3-3 binding, with only a modest reduction compared to WT BRAF (Fig. 4B). In addition, S729 phosphorylation was not significantly affected by the A2 (or R2) mutation (Fig. 4C), indicating that the loss of RNA binding is unlikely to result from altered C-terminal regulatory phosphorylation and impaired 14-3-3 recruitment.

We next examined the effect of these mutations on kinase activity by itself. Using the constitutively active BRAF V600E, we found that disruption of the basic surface (3E) did not affect MEK1/2 phosphorylation, in contrast to a kinase-impaired BRAF deletion mutant lacking part of the C-terminal domain (Δ718-766) (Fig. 4D). These results suggest that disruption of the basic surface does not compromise the intrinsic kinase activity of BRAF. To further determine whether the A2 mutation affects the intrinsic catalytic activity of BRAF, we performed *in vitro* kinase assays using immunoprecipitated proteins. BRAF V600E as well as BRAF V600E A2 and BRAF V600E Δ718-766 were immunoprecipitated to comparable levels from transfected HEK293T cells and incubated with recombinant inactive MEK1 as a substrate. The BRAF V600E A2 mutant phosphorylated MEK1 as efficiently as BRAF V600E (Fig. 4E), indicating that disruption of the RNA-binding interface does not impair the intrinsic kinase activity of BRAF. The kinase-impaired BRAF V600E Δ718-766 markedly diminished MEK1 phosphorylation under the same assay conditions (Fig. 4E). Together with its preserved 14-3-3 association, these data indicate that the A2 mutant is a separation-of-function mutant that selectively disrupts RNA binding while maintaining the catalytic properties of the kinase.

If the A2 mutation represents a separation-of-function allele that selectively disrupts RNA binding while preserving other BRAF activities, then restoring RNA proximity should rescue its signaling defect. To test this, we developed a synthetic RNA recruitment system based on the high-affinity interaction between the bacteriophage MS2 coat protein (MCP) and its cognate MS2 RNA stem-loop (Peabody 1993). MCP was fused to the RNA-binding-deficient BRAF A2 mutant and co-expressed with a reporter RNA carrying an MS2 binding site downstream of the luciferase ORF, thereby recruiting two BRAF molecules to an artificial RNA, independently of the endogenous RNA-binding interface (Fig. 4F). Expression of the MS2-containing reporter (Luc-MS2), but not a control reporter carrying an unrelated 212-nt sequence (Luc-ctl), increased MEK1/2 phosphorylation by 2.1-fold in cells expressing BRAF A2-MCP (Fig. 4G). In contrast, in cells expressing wild-type BRAF-MCP, RNA recruitment produced a weaker 1.4-fold increase in MEK1/2 phosphorylation (Fig. 4G), consistent with the intact endogenous RNA-binding interface in WT BRAF. These results demonstrate that enforced RNA-mediated dimerization of BRAF restores signaling by the RNA-binding-deficient BRAF A2 mutant, providing evidence that RNA association is involved in BRAF signaling.

## DISCUSSION

Our data establish RNA as a direct regulator of BRAF signaling and reveal an additional mechanism controlling RAF activity. BRAF binds RNA in cells, with RNA interaction increasing upon MAPK pathway activation, while depletion of intracellular RNA reduces BRAF:CRAF proximity and MEK1/2 phosphorylation. This mechanism is distinct from the described role of snoRNAs *SNORD50A/B*, which bind KRAS (Siprashvili et al. 2016) and antagonize its SNARE-mediated trafficking to the plasma membrane (Che et al. 2019). Our study identifies an RNA-BRAF interaction as a previously unrecognized layer of RNA-mediated regulation of the MAPK pathway, acting downstream of RAS and upstream of MEK.

The identification of a basic surface within the BRAF kinase domain that mediates RNA binding provides a molecular basis for this regulation. Importantly, the K723A separation-of-function mutant uncouples RNA binding from the intrinsic catalytic activity of BRAF and from its association with 14-3-3, yet impairs MAPK signaling. Conversely, artificially recruiting this mutant to an RNA scaffold restores signaling activity. Together, these results support a model in which RNA promotes the productive assembly or stabilization of RAF complexes rather than simply modulating BRAF catalytic activity. RNA may therefore function as a molecular scaffold increasing the local concentration or appropriate folding of RAF molecules required for efficient kinase activation.

The repertoire of endogenous RNAs interacting with BRAF is likely to vary with the cellular transcriptome. Cancer cell type-specific RNA transcriptomes may influence the magnitude and dynamics of RAF-MAPK signaling and thereby contribute to the behavior of MAPK-driven tumors. In addition, resistance to BRAF V600E inhibitors can involve increased RAF complex formation (Boussemart et al. 2016, 2014), raising the possibility that RNAs enriched in resistant cells contribute to this process by promoting RAF assembly. Overall, determining whether RNA-dependent regulation of RAF signaling varies across oncogenic RAS/BRAF-driven tumors and between therapy-sensitive and therapy-resistant states will require identification and characterization of the endogenous RNAs that interact with BRAF in these contexts. This could also pave the way for evaluating RNA-BRAF interactions in tumor biopsies as prognostic or predictive biomarkers of therapeutic response.

## KEY RESOURCES TABLE

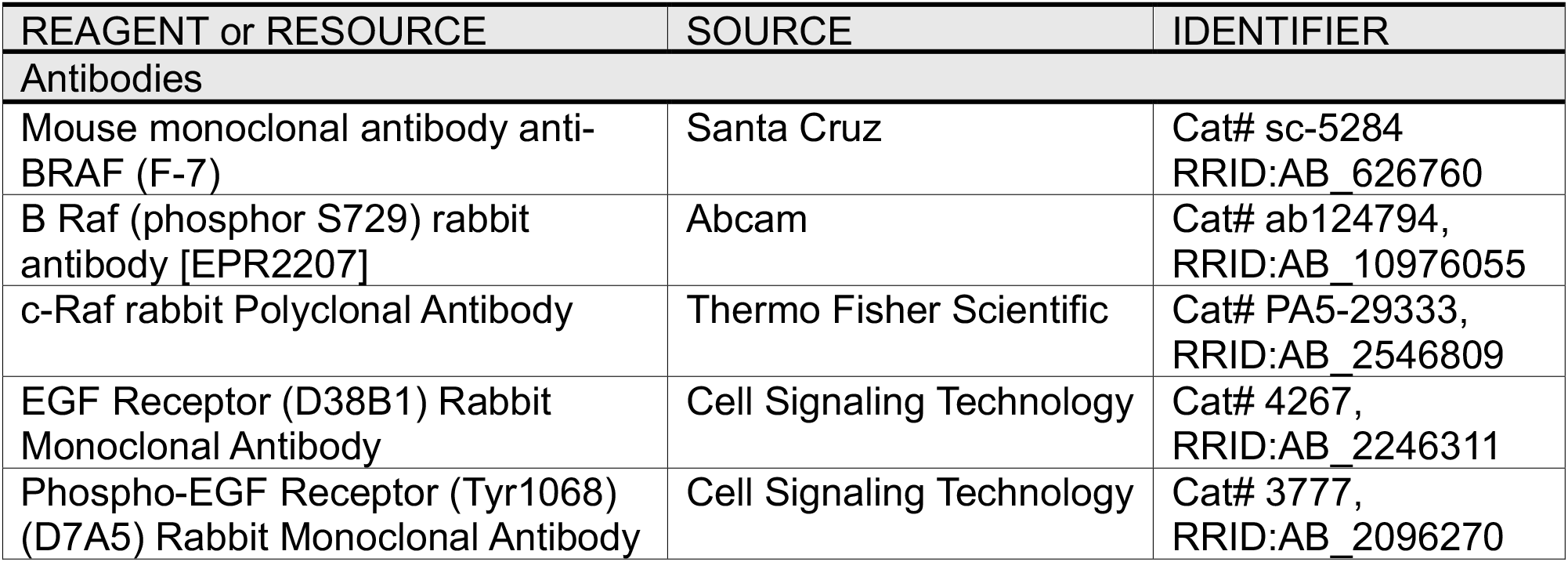
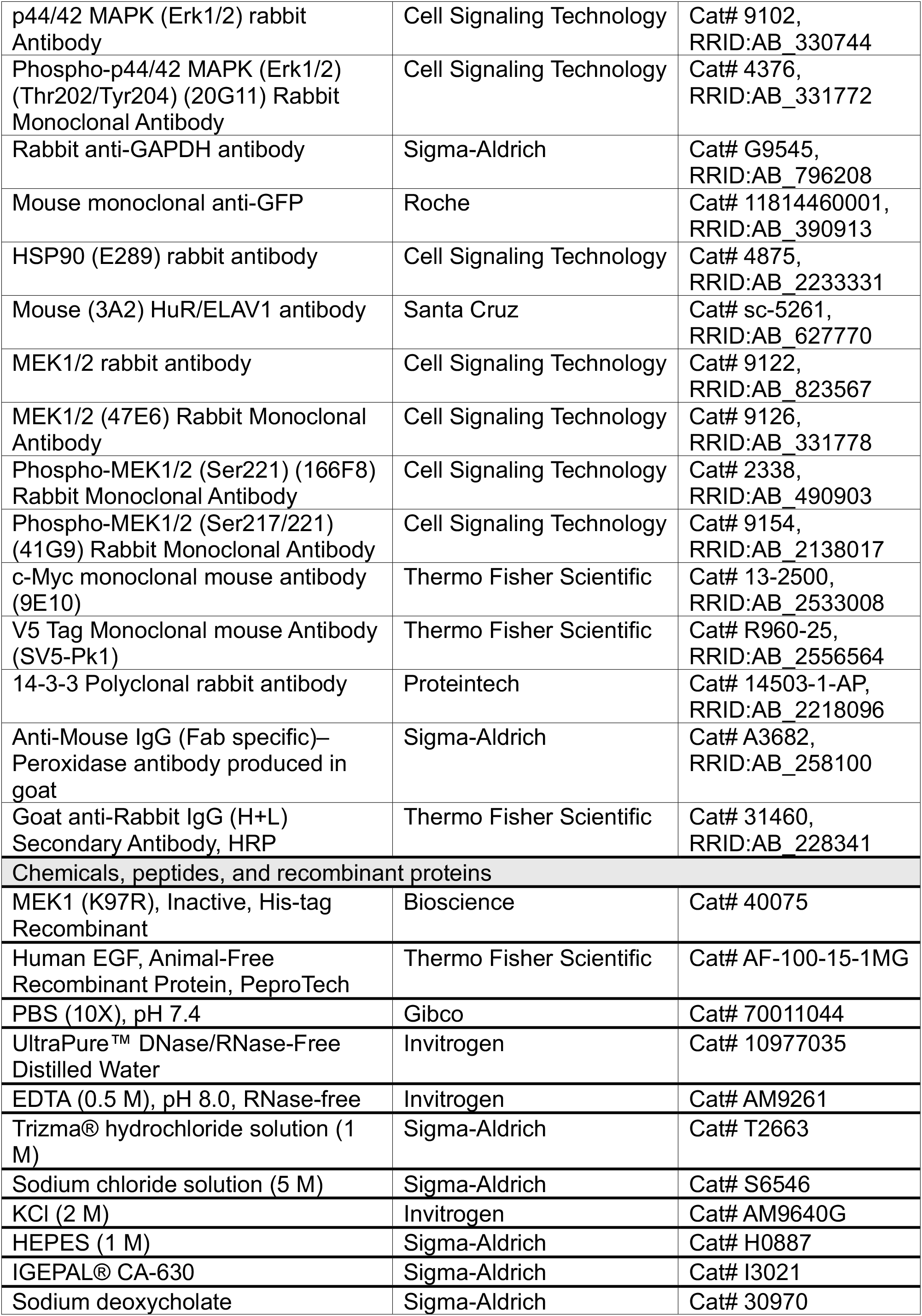
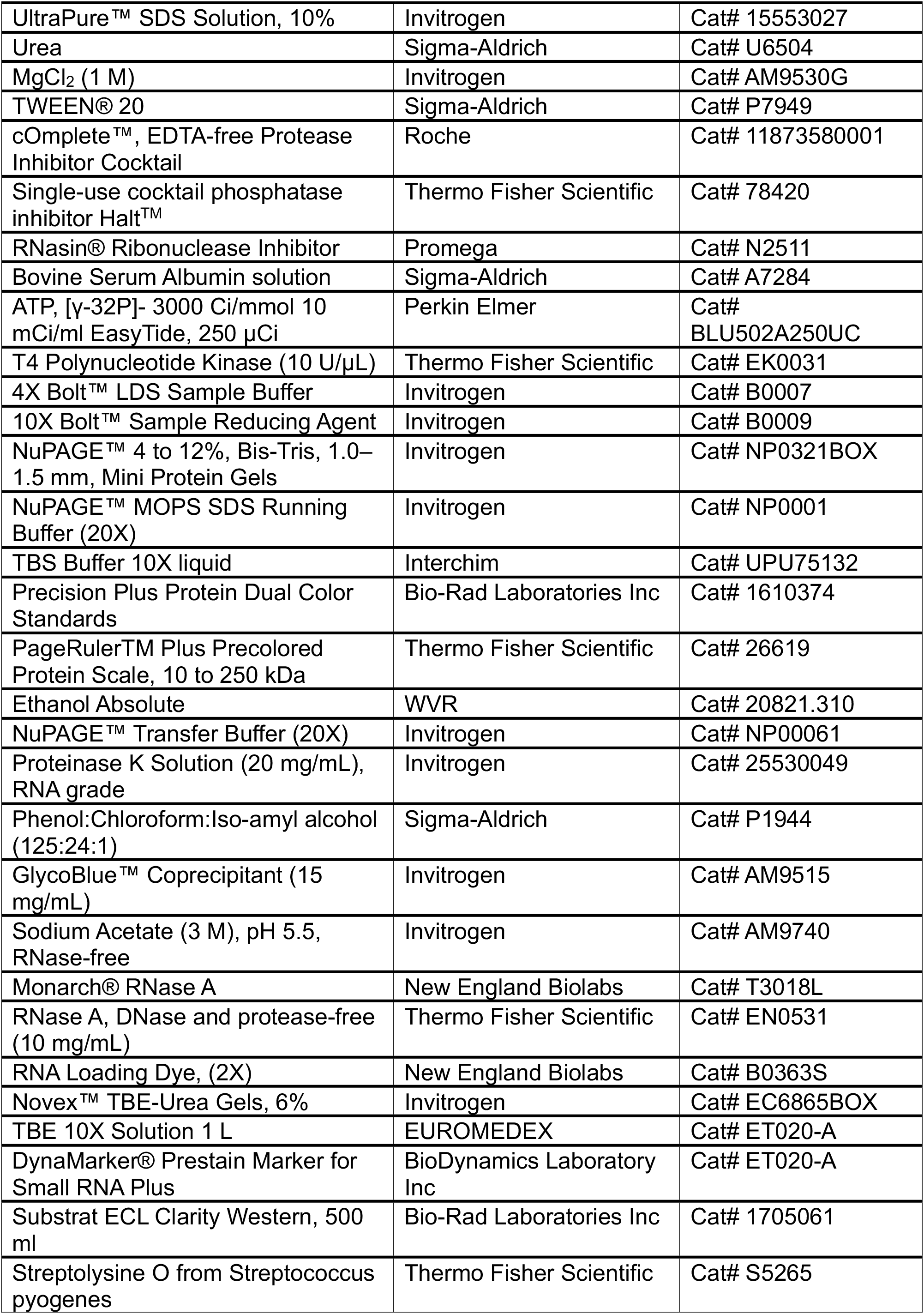
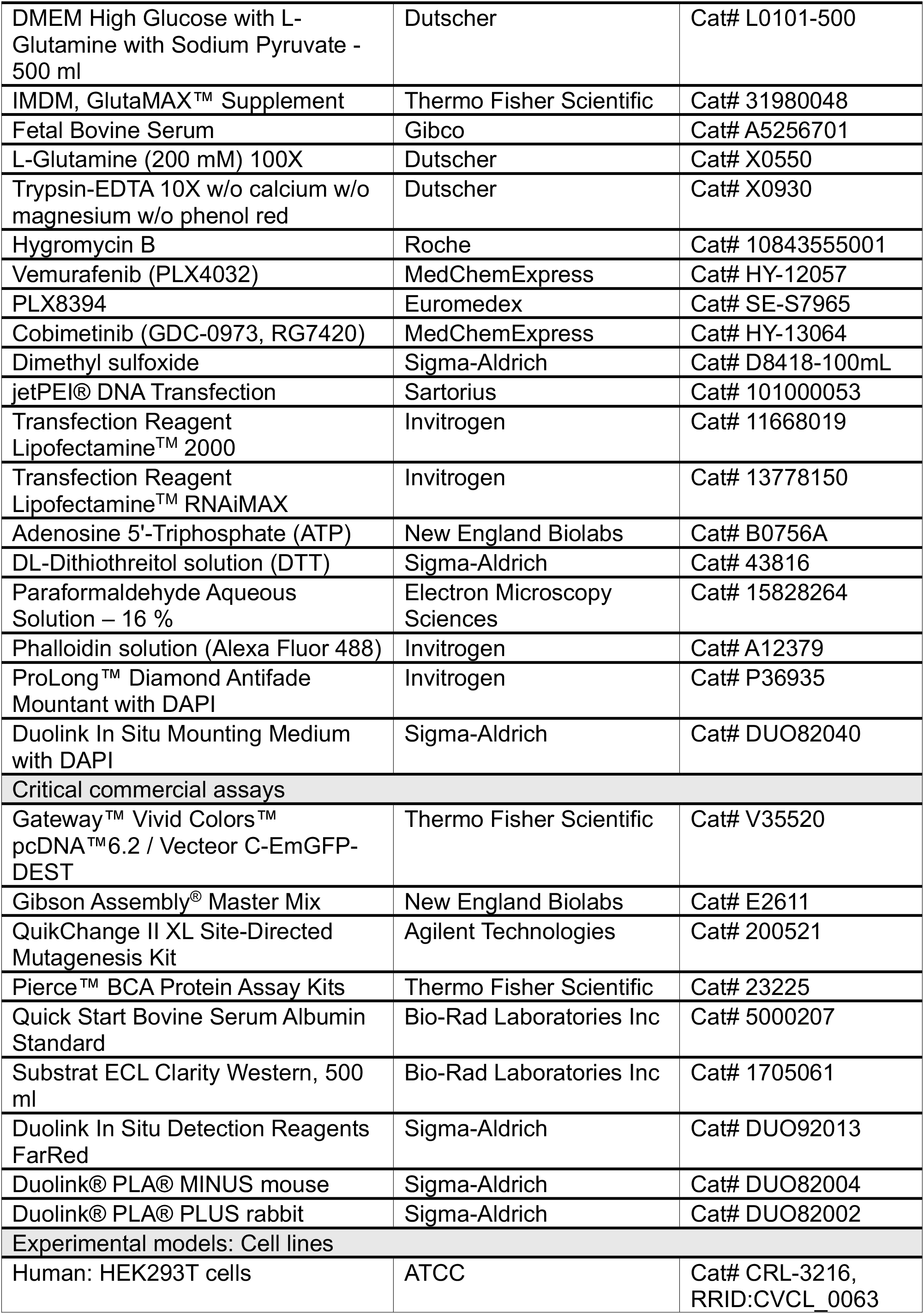
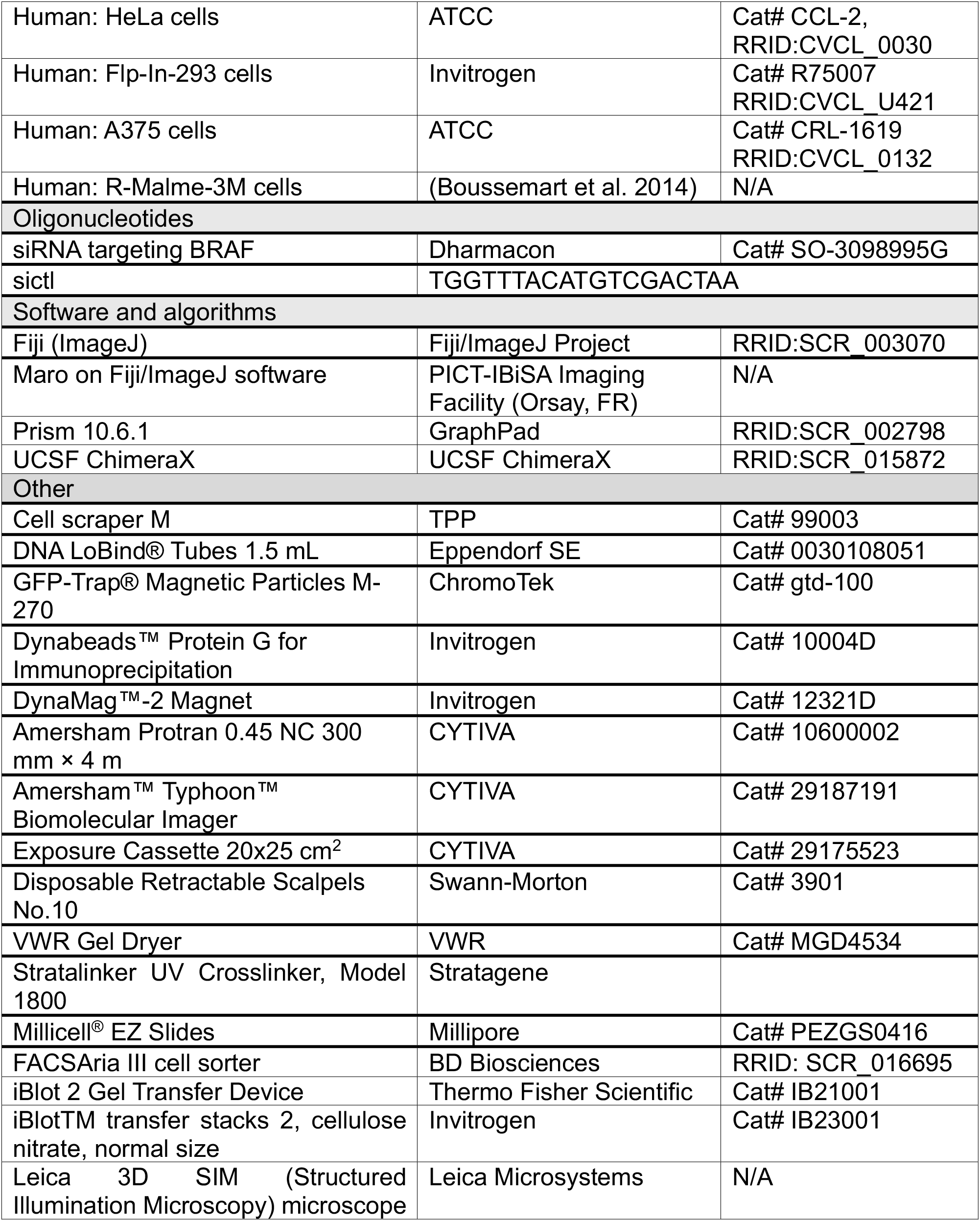

## METHODS

### Cell culture

HEK293T, HeLa and A375 cell lines were purchased from the American Type Culture Collection (ATCC). Flp-In™-293 (Invitrogen™, #R75007) cell line was purchased from Invitrogen. R-Malme-3M were previously generated in the laboratory as described in a previous study (Boussemart et al. 2014). HEK293T, HeLa and A375 and Flp-In™-293 cells were grown in high glucose DMEM (Dutscher, #L0101-500) supplemented with 10 % FBS (Gibco™, #A5256701) and 2 mM L-glutamine (Dutscher, #X0550-100). R-Malme-3M cells were grown in high glucose IMDM (Gibco, #12589059) supplemented with 20 % FBS, and treated with 1 µM vemurafenib (MedChemExpress, #HY-12057). All cells were grown in 5 % CO_2_ humidified incubator at 37°C. Flp-In™-293 were used to generate a cell line expressing BRAF-V5, according to manufacturers’ instructions. The established cell line was cultured in high glucose DMEM medium with 10 % FBS and 2 mM L-glutamine supplemented with 100 µg/mL hygromycin B (Thermo Fisher Scientific, #10687010). Trypsin (Dutscher, #X0930) was used to pass cell and harvest cells, when not scrapped.

To synchronize the cells, HeLa cells were incubated for 16 h in high glucose DMEM supplemented with only 2 mM L-glutamine. The cells were washed with PBS before the medium was replaced. After 16 h of starvation, the cells were stimulated with 0.1 µg/mL EGF (Thermo Fisher Scientific, #AF-100-15-1MG) for 5 minutes. When indicated, cells were also permeabilized using Streptolysine O (SLO ; 150 U/mL) for 10 min at 37°C and treated with RNAse A (1 mg/mL) for 10 min at 37°C before being stimulated with EGF and subsequently harvested. SLO was preactivated with 20 mM DTT for 20 min at 37°C. Various MAPK pathway inhibitors were used to treat BRAF V600E expressing cells for 1 h, it included vemurafenib/PLX4032 (MedChemExpress, #HY-12057) at 1 µM, PLX8394 (Euromedex, #SE-S7965) at 1 µM and cobimetinib (MedChemExpress, #HY-13064) at 0.1 µM.

Cells were regularly tested for mycoplasma and authenticated by short tandem repeat (STR) profiling.

### Plasmid cloning and mutagenesis

NRAS-myc WT and Q61K-mutated were kindly given from Eychène team (Institut Curie). MS2-HB (Addgene, #35573) and Tagged-Cap (Addgene, #35572) plasmids were purchased from Addgene. BRAF-GFP and CRAF-GFP were designed and cloned using the Gateway™ cloning technology Gateway™ Vivid Colors™ pcDNA™6.2 / Vecteor C-EmGFP-DEST following the manufacturers’ instructions (Thermo Fisher Scientific, #V35520). Site-directed mutagenesis was performed on BRAF-GFP coding pcDNA6.2 using QuikChange II XL kit (Agilent Technologies, #200521) and primers specifically introducing each following mutations: S365A, R509H, V600E, R719A, K723A, K723R, R726A, S729A or the combinations of three substitutions: R719E-K723E-R726E and R719A-K723A-R726A. Gibson assembly (NEB, # E2611) strategy was used on BRAF-GFP pcDNA6.2 to fuse BRAF-GFP to MCP protein, according to manufacturers’ instructions. MCP cDNA was generated by PCR on the MS2-HB plasmid. Gibson strategy was also used to remove two of the three MS2 stem-loops initially present in the Tagged Cap plasmid.

### Transfections

Cells were plated 24 h before siRNA and plasmid transfections. HEK293T cells were transfected using JetPei (Sartorius, #101000053), while Flp-In™-293 according to manufacturers’ instructions. At 48 h post-transfection, cells were harvested for further analysis. The siRNAs were purchased from Dharmacon (ON-TARGETplus technology). siRNA transfections were performed at 10 nM using Lipofectamine RNAiMAX (Invitrogen™, #13778150) according to manufacturer’s instructions.

### Protein extracts, SDS-PAGE and western blot

For western blot performed on BRAF-GFP transfected HEK293T cells, cells were harvested after 48 h of transfection and 800,000 high-GFP-expressing cells were sorted using a FACSAria III (BD Biosciences) cell sorter at the CYTPIC - cytometry facility in Orsay. Whole cell lysates were prepared using RIPA buffer (50 mM Tris-HCl pH 7.4, 100 mM NaCl, 1 % Nonidet P-40, 0.1 % SDS, 0.5 % Sodium deoxycholate) supplemented with 40 U/mL RNasin (Promega, #N2511), 1X protease inhibitor cocktail (Roche, #11873580001) and 1X phosphatase inhibitor cocktail (Thermo Scientific™, #78420). Lysates were cleared via centrifugation at 18,000 g for 20 minutes at 4°C. Cell lysates were quantified for protein concentration using a bicinchoninic acid (BCA) protein assay kit (Thermo Scientific™, #23225). Protein samples were resolved on NuPAGE 4–12 % Bis-Tris gels (Invitrogen™) with 1X MOPS buffer (Invitrogen™, #NP0001) and then transferred to 0.45 μm nitrocellulose membrane (Invitrogen™, IB23001) using the dry iBlot 2 Gel Transfer Device (Thermo Fisher Scientific, #IB21001). After saturation in Tris-buffered saline buffer containing 0.01% Tween-20 (TBST) and supplemented with 5 % powdered milk, the membranes were incubated with primary antibodies either for 1 hour at room temperature or overnight at 4 °C with agitations, followed by three TBST washes of 5 minutes each, and incubation of the membranes with peroxidase (HRP)-conjugated secondary antibodies diluted in 5 % milk TBST for 1 hour at room temperature. After three TBST washes of 5 minutes each, the membranes were incubated in ECL solution (Bio-Rad, #1705061) for signal detection. The antibodies used are presented in the key resources table. The intensity of phospho-proteins and total proteins on blots were quantified with ImageJ using “Gel” tool. Statistical tests were performed on phospho/total ratios and ratio fold change compared to a control condition fixed to 1 is indicated below each gel.

### Co-Immunoprecipitation (Co-IP)

Cells were cultured on 10 cm dishes and after 24 h, they were transfected for 48 h. Cells were harvested and lysed for 30 minutes on ice in 1 mL Co-IP lysis buffer (10 mM HEPES pH7.4, 10 mM KCl, 150 mM NaCl, 0.1 % Nonidet P-40), supplemented with 40 U/mL RNasin (Promega, #N2511), 1X protease inhibitor cocktail (Roche, #11873580001) and 1X phosphatase inhibitor cocktail (Thermo Scientific™, #78420). Lysates were cleared via centrifugation at 18,000 g for 15 minutes at 4°C. Cell lysates were quantified for protein concentration using a bicinchoninic acid (BCA) protein assay kit (Thermo Scientific™, #23225). Cell lysates were incubated with 30 µL GFP-Trap® Magnetic Particles M-270 (ChromoTek, #gtd-100) overnight at 4°C while constantly rotating. The immunocomplexes were washed four times with ice-cold wash buffer (10 mM HEPES pH7.4, 10 mM KCl, 150 mM NaCl, 0.1 % Nonidet P-40, 1 mM EDTA) then they were eluted in 1X loading buffer at 80°C for 5 minutes and separated by SDS-PAGE (see western blot method for following steps).

### Kinase Assay

The protocol was adapted from the BRAF kinase assay kindly given by Lavoie’s lab (personal communication). Cells were cultured on 10 cm dishes and after 24 h, they were transfected for 48 h. Cells were harvested and lysed for 15 minutes on ice in 1 mL Igepal lysis buffer (50 mM Tris-HCl pH7.5, 150 mM NaCl, 1 % Nonidet P-40, 1 mM EDTA, 10 % glycerol), supplemented with 40 U/mL RNasin (Promega, #N2511), 1X protease inhibitor cocktail (Roche, #11873580001) and 1X phosphatase inhibitor cocktail (Thermo Scientific™, #78420). Lysates were cleared via centrifugation at 18,000 g for 10 minutes at 4°C. Cell lysates were incubated with 50 µL GFP-Trap® Magnetic Particles M-270 (ChromoTek, #gtd-100) overnight at 4°C while constantly rotating. 1/50 of the immunoprecipitated protein was incubated for 30 minutes at 30°C in KBA buffer (50 mM HEPES pH7.5, 10 mM MgCl_2_, 1 mM EDTA) with 200 µM ATP (NEB, #B0756A), 9 ng/µL inactive MEK1 (Bioscience, #40075) and 2 mM DTT. Phosphorylation of MEK1 was then assessed by western blot.

### Proximity Ligation Assay (PLA)

The experiment was performed using Duolink In situ Detection Reagents FarRed according to the manufacturer’s instructions (Sigma-Aldrich, #DUO92013). HeLa cells were seeded on Millicell^®^ EZ Slides (Millipore, # PEZGS0416) 24 h before starvation and EGF stimulation. Cells were permeabilized with Tween 0.2 % in PBS for 10 minutes at room temperature and then RNAse A treatments were performed 2 minutes at 37°C. Next, cells were fixed with 4 % PFA (Electron Microscopy Sciences, #15828264) in PBS for 10 minutes at room temperature. Cells were then washed with PBS and blocked using the kit blocking solution for 20 minutes at 37°C. Cells were incubated in antibodies mix for 1 h at 37°C, then washed twice with kit Buffer A. The antibodies used are presented in the key resources table. Then cells were incubated 1 h at 37°C with Duolink® PLA® MINUS and Duolink® PLA® PLUS probes and washed twice with kit Buffer A. A ligation solution was then added to ligate the complementary oligonucleotides of the two probes (PLUS and MINUS), by incubating the coverslips 30 minutes at 37°C. After, two washes with kit Buffer A, polymerase-mediated amplification by rolling circle replication was performed using the kit polymerase solution incubated 1 h 30 min at 37°C. After two washes with kit Buffer B, and one with 1/100-diluted Buffer B, cytoplasms were stained using 1X phalloidin solution (Alexa Fluor 488; Invitrogen™, #A12379) for 20 minutes at room temperature. After PBS washes, the coverslips were finally mounted in ProLong™ Diamond Antifade Mountant with DAPI (Invitrogen™, #P36935). Images of each coverslip were acquired using Leica 3D SIM (Structured Illumination Microscopy) microscope at the PICT-IBiSA Imaging Facility in Orsay and PLA dots were counted using the semi-automatic macro on ImageJ software, created by the PICT-IBiSA Imaging Facility.

### Crosslinking and immunoprecipitation

Cells were cultured on 10 or 15 cm dishes until 70-80 % of confluence or until 48 h of transfection. The medium was removed, and the cells were washed with ice-cold PBS and irradiated with UV-C light at 254 nm. Cells were then lysed in 1 mL RIPA lysis buffer (50 mM Tris-HCl pH 7.4, 100 mM NaCl, 1 % Nonidet P-40, 0.1 % SDS, 0.5 % Sodium deoxycholate), supplemented with 40 U/mL RNasin (Promega, #N2511) and protease inhibitor cocktail (Roche, #11873580001). Next, lysates were cleared via centrifugation at 18,000 g for 20 minutes at 4°C, and were quantified for protein concentration using a bicinchoninic acid (BCA) protein assay kit (Thermo Scientific™, #23225). Then lysates were treated with 10 µg/mL RNAse A (Thermo Scientific™, #EN0531), using approximatively 10 µg RNAse A for 1 mg of proteins and incubated at 37°C for 5 minutes at 800 rpm. 2 to 5 % of the lysate was used as input material to quantify total protein concentration in western blot analysis. The rest of the lysate was incubated with BRAF antibody-conjugated beads or GFP-Trap® Magnetic Particles M-270 (ChromoTek, #gtd-100) overnight at 4°C while constantly rotating. For endogenous BRAF immunoprecipitation, 6 µg of BRAF antibody (Santa Cruz, #sc-5284) were coupled 4 h at 4°C to 50 µl Dynabeads Protein G (Invitrogen™, #10004D) while constantly rotating and wash prior mixing with the samples. For GFP-tagged protein immunoprecipitation, 40 µL of GFP-Trap® Magnetic Particles M-270 were directly used. The immunocomplexes were washed twice with RIPA-S buffer (50 mM Tris-HCl pH 7.4, 1 M NaCl, 1 mM EDTA, 1 % Nonidet P-40, 0.1 % SDS, 0.5 % Sodium deoxycholate) and once with PNK buffer (20 mM Tris-HCl pH 7.4, 10 mM MgCl_2_, 0.2 % Tween 20). After washes, 10 % of the immunocomplexes was used to validate the correct immunoprecipitation by western blot analysis. Then, crosslinked nucleic acids were radiolabeled with ATP, [γ-32P] (PerkinElmer, #NEG002A250UC) by T4 polynucleotide kinase (in PNK and B buffers; Thermo Scientific™, #EK0031) at 37°C for 15 minutes at 800 rpm. After two washes with RIPA-S buffer and one wash with PNK buffer, protein-RNA complexes were eluted in 1X loading buffer at 80°C for 5 minutes and separated by SDS-PAGE. Then the gel was dried, and complexes were exposed to a phosphorimaging screen and scanned with Amersham™ Typhoon™ Biomolecular Imager (CYTIVA). The files were processed with ImageJ software. When indicated, RNA-proteins complexes were transferred onto nitrocellulose membrane 0.45 µm (Amersham, #10600002), instead of direct gel drying. The radiolabeled complexes were similarly visualized by exposure to a phosphorimager screen. The nitrocellulose membrane was cut out from the suspected molecular weight of the radiolabeled protein of interest to 60 kDa above. Nitrocellulose pieces were treated with proteinase K (1 mg/mL; Thermo Scientific, #EO0491) in PK buffer (100 mM Tris-HCl pH 7.4, 50 mM NaCl, 10 mM EDTA), once without urea and then once with urea 7 M, for 20 minutes at 37°C each. Radioactively-labelled nucleic acids were purified with phenol/chloroform/isoamylic alcohol 125:24:1 (Sigma, #P1944) and then precipitated overnight by adding Glycoblue (Thermofischer, #AM9515), sodium acetate and 100 % ethanol. After centrifugation and washes (80 % ethanol), the recovered nucleic acids pellets were resuspended in water and divided in two to assess RNase degradation. Untreated and RNAse A-treated RNAs were separated using denaturing polyacrylamide gel (TBE-Urea gel, 6 %), which was dried and exposed to a phosphorimaging screen. Complete RNase-degradation confirms the nature of the nucleic acid recovered.

### Orthogonal Organic Phase Separation (OOPS)

Cells were cultured on 10 or 15 cm dishes until 70-80 % of confluence or until 48 h of transfection. The medium was removed, and the cells were washed with ice-cold PBS and irradiated with UV-C light at 254 nm. Cells were then lysed in Trizol (Invitrogen, #15596018) and lysates were subjected to acid guanidinium thiocyanate–phenol– chloroform (AGPC) phase separation according to the manufacturer’s instructions. Following centrifugation, the aqueous phase containing RNA and the organic phase containing proteins were discarded, while the interphase, enriched in covalently crosslinked RNA–protein complexes, was retained. RBPs were recovered after RNase treatment and their presence in the interface was assessed by western Blot. RNAse treatment performed during lysis was used as negative control and HuR was used as a positive control.

### Quantification and statistical analysis

Statistical analyses were performed on GraphPad Prism (Version 10.6.1) and statistical significance defined as a P value < 0.05 was determined by two-tailed unpaired or paired t-test, one-way or 2-way ANOVA, as indicated in the figure legends. All data in this study are presented as the mean standard deviation or mean standard error.

## ACKNOWLEDGEMENTS

This work was supported by grants from Institut Curie, Gustave Roussy, INSERM, CNRS, Université Paris-Saclay, Equipe labellisée Ligue Nationale Contre le Cancer (LNCC) (to SV), Institut national du cancer (INCA) (to SV and CR) and la Fondation du Crédit Mutuel (to CR). ALB was successively supported by pre-doctoral fellowships from Ecole Polytechnique and Fondation ARC pour la recherche sur le cancer. SM was successively supported by pre-doctoral fellowships from the Ministère de l’Enseignement Supérieur et de la Recherche (MESR) and Fondation ARC pour la recherche sur le cancer. We thank Martin Dutertre, Bruno Galy, Matthias Hentze and Marc Therrien for helpful discussions. We thank Sylvain Martineau and Vanessa Caicedo Lemos for their help with the cloning and permeabilization/western blot experiments, respectively. We are grateful to Charlène Lasgi from the Flow Cytometry platform of Institut Curie and Laetitia Besse from the Multimodal Imaging Center Imaging Facility of the Institut Curie (CNRS UAR2016/INSERM US43/Institut Curie/Université Paris-Saclay) for their support with the experiments.

## AUTHOR INFORMATION

Conceptualization: ALB, SM, LF, PU, SV

Data curation: ALB, PU, SV

Formal analysis: ALB, SM, LF, TS, IG, VQ, JS, JA, RG, CR, PU, SV

Funding acquisition: CR, SV

Investigation: ALB, SM, LF, TS, IG, VQ, JS, JA, RG, PU, SV

Methodology : ALB, SM, LF, TS, IG, VQ, JS, JA, RG, PU, SV

Project administration : SV

Ressources : SV, CR

Supervision : SV, PU, CR

Writing original draft : SV with the help of ALB and PU

Writing – review & editing : ALB, SM, LF, PU, SV

## DATA AVAILABILITY

All data are available in the main text or the supplementary materials.

## FIGURE LEGENDS FOR SUPPLEMENTARY FIGURES

**Figure S1.**
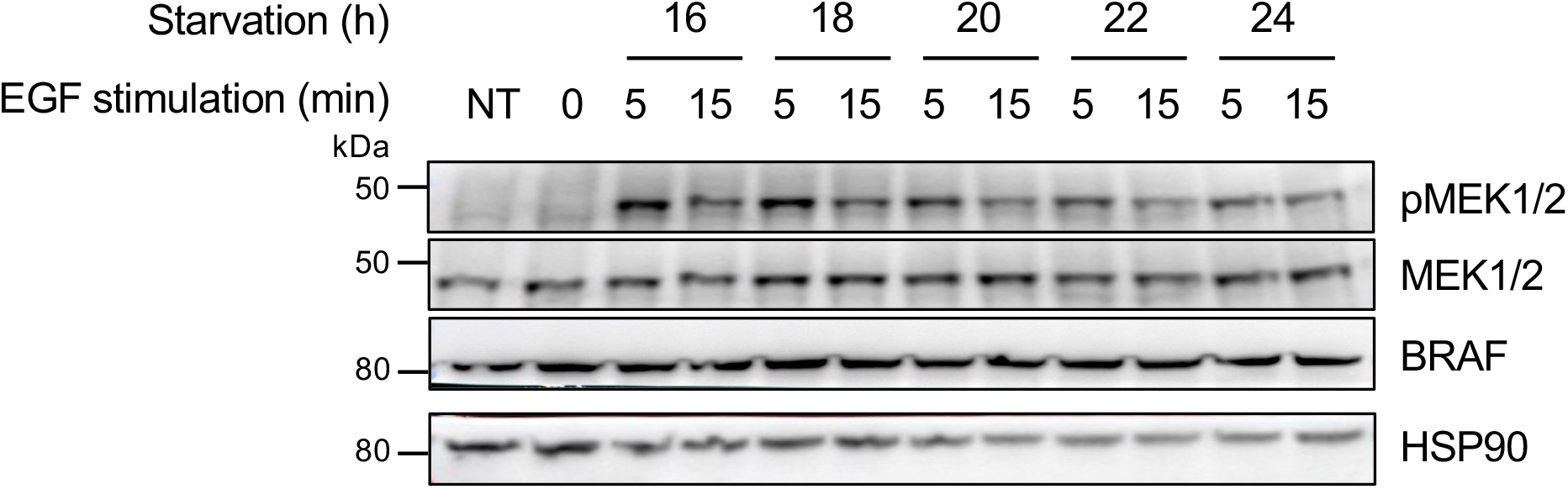
Immunoblot analysis of HeLa cells that were serum-starved for 16 to 24 h and stimulated with EGF for 5 and 15 minutes. HSP90 was used as a loading control. NT: no treatment.

**Figure S2.**
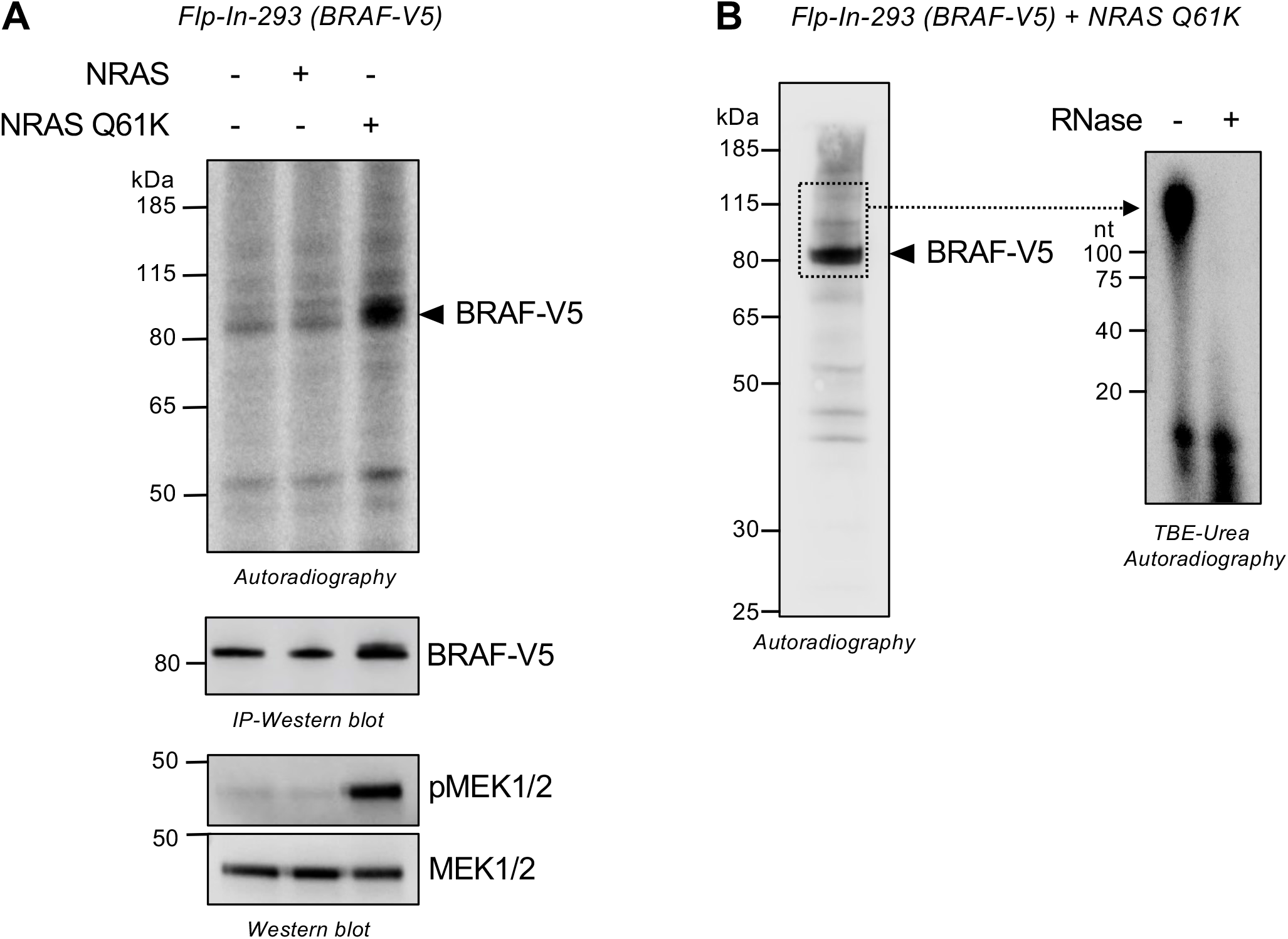
**(A)** CLIP of stably expressing V5-tagged BRAF in Flp-In-293 cells (n = 3 biological replicates). Upper panel: Autoradiography of BRAF-RNA complexes from UV-irradiated cells transfected with NRAS or NRAS Q61K. A prominent band corresponding to the expected molecular weight of BRAF is indicated by a black arrow. Middle panel: Immunoblot of BRAF immunoprecipitates under the same experimental conditions shown in the upper panel. Lower panel: Immunoblot performed under the same experimental conditions. **(B)** CLIP of stably expressing V5-tagged BRAF in Flp-In-293 cells (n = 3 biological replicates). Left panel: Autoradiography of BRAF-RNA complexes from UV-irradiated cells. Right panel: Autoradiography of extracted nucleic acids. Radioactively labeled nucleic acids were extracted from membrane sections corresponding to the dashed squares in the left panel and run on a denaturing TBE-Urea gel. RNase digestion (+) was performed on half of each sample to confirm the RNA nature of the detected signal.

**Figure S3.**
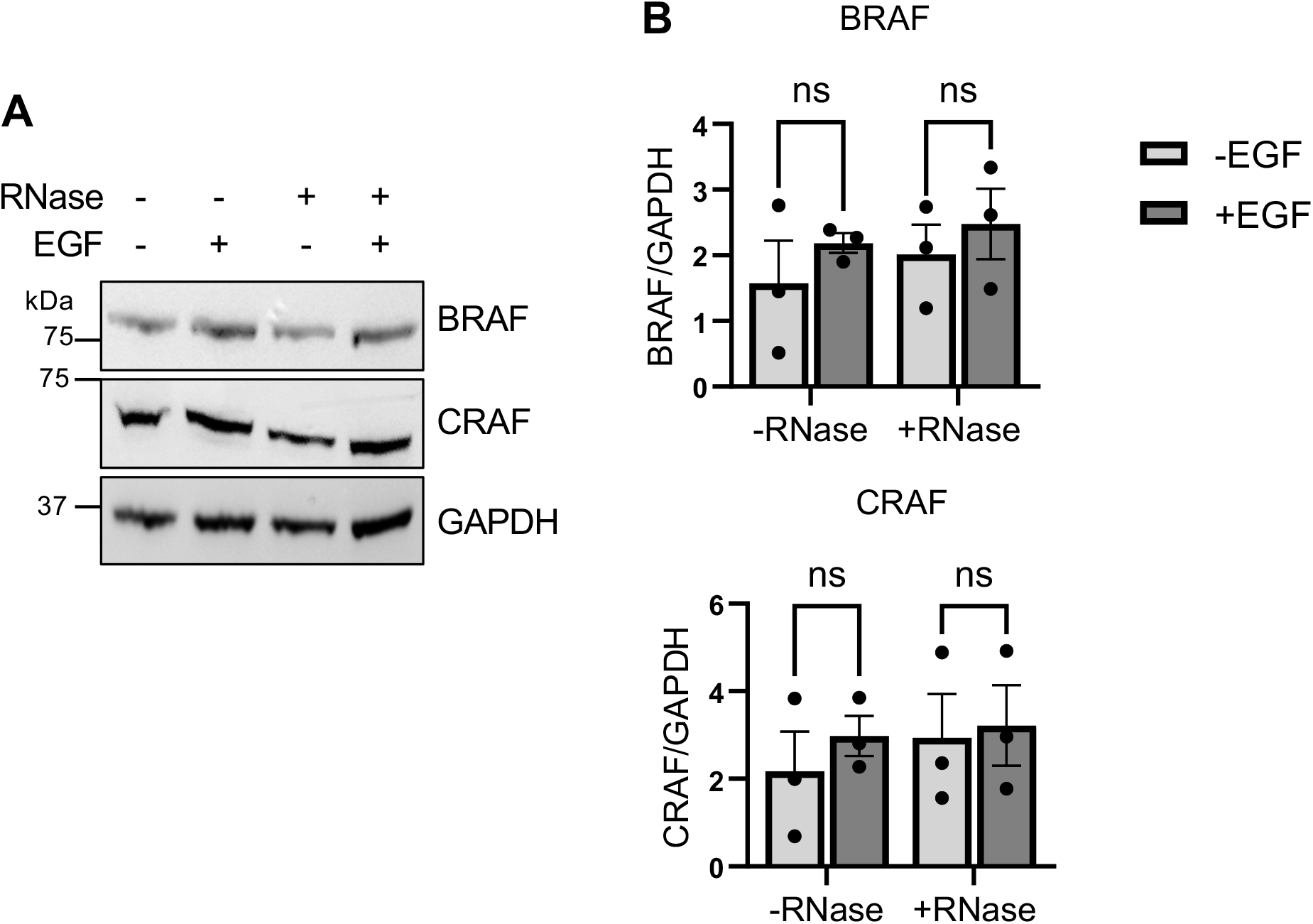
**(A**) Immunoblot analysis of BRAF and CRAF protein levels in serum-starved (16 h) HeLa cells stimulated with EGF for 5 min (+) or left unstimulated (-), permeabilized with Tween-20 and subsequently treated with (+) or without (-) RNase A. GAPDH was used as a loading control. **(B)** Histograms showing BRAF (upper) or CRAF (lower) protein levels, normalized to GAPDH, under the indicated conditions. P values were determined using ordinary 2-way ANOVA followed by Sidak’s multiple-comparisons (BRAF: -RNase : –EGF vs +EGF, *P* = 0.6366 ; +RNase: -EGF vs +EGF, *P* = 0.7575 / CRAF: -RNase : –EGF vs +EGF, *P* = 0.7705 ; +RNase: -EGF vs +EGF, *P* = 0.9680).

**Figure S4.**
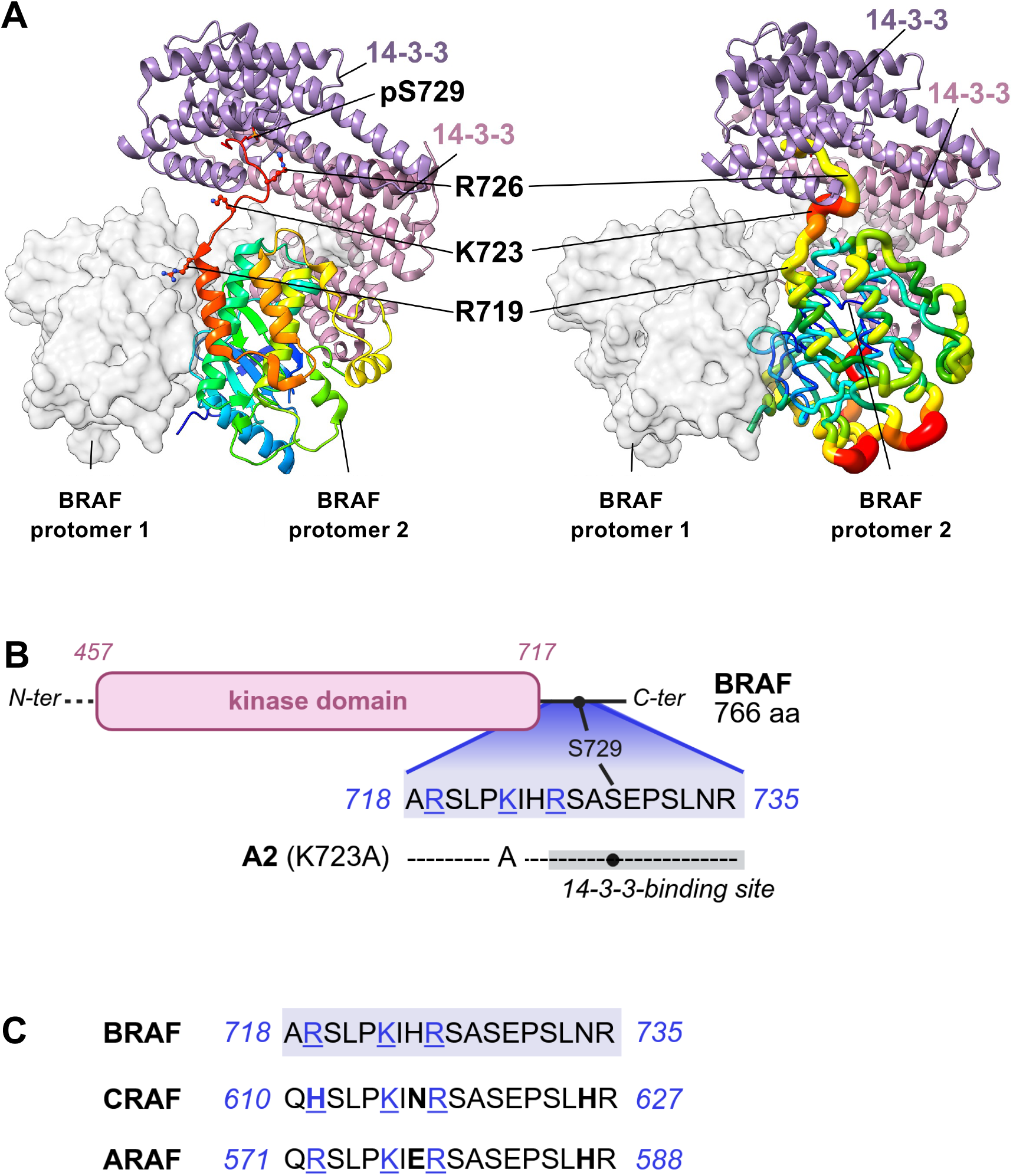
**(A)** Distribution of crystallographic B-factors in BRAF within the BRAF-14-3-3 complex. The crystal structure of the apo BRAF dimer bound to a 14-3-3 dimer (PDB 6XAG) is shown. Left panel, the complex is represented as in Figure 4, with BRAF protomer 1 shown as a white molecular surface, BRAF protomer 2 as a rainbow-colored cartoon and the two 14-3-3 protomers in purple and pink. The C-terminal BRAF residues R719, K723, R726, and pS729 are indicated. In the right panel, BRAF protomer 2 is displayed in a representation in which tube thickness and rainbow coloration reflect the crystallographic B-factor values, ranging from lower B-factors in blue/cyan to higher B-factors in yellow/orange/red. This representation highlights differences in local structural mobility within BRAF, particularly at K723, which displays one of the highest B-factor values, consistent with increased local mobility and a lack of stabilizing contacts with surrounding structural elements. BRAF protomer 1 and the 14-3-3 dimer are shown for structural context. Molecular graphics and structural analyses were performed using UCSF ChimeraX (Pettersen et al. 2021). **(B)** Schematic representation of the 14-3-3-binding site in the vicinity of the A2 mutation. S729 phosphosite responsible for 14-3-3 interaction is indicated with a black dot. **(C)** Sequence alignment of the region encompassing the A2 mutation in all three RAF isoforms.

## BIBLIOGRAPHY

Abe, Yohei, Eric R. Kofman, Maria Almeida, et al. 2023. ‘RANK Ligand Converts the NCoR/HDAC3 Co-Repressor to a PGC1β- and RNA-Dependent Co-Activator of Osteoclast Gene Expression’. Molecular Cell 83 (19): 3421–3437.e11. 10.1016/j.molcel.2023.08.029.

Baltz, Alexander G., Mathias Munschauer, Björn Schwanhäusser, et al. 2012. ‘The mRNA-Bound Proteome and Its Global Occupancy Profile on Protein-Coding Transcripts’. Molecular Cell 46 (5): 674–90. 10.1016/j.molcel.2012.05.021.

Beckmann, Benedikt M., Alfredo Castello, and Jan Medenbach. 2016. ‘The Expanding Universe of Ribonucleoproteins: Of Novel RNA-Binding Proteins and Unconventional Interactions’. Pflügers Archiv - European Journal of Physiology, ahead of print. 10.1007/s00424-016-1819-4.

Boussemart, Lise, Isabelle Girault, Hélène Malka-Mahieu, et al. 2016. ‘Secondary Tumors Arising in Patients Undergoing BRAF Inhibitor Therapy Exhibit Increased BRAF-CRAF Heterodimerization’. Cancer Research 76 (6): 1476–84. 10.1158/0008-5472.CAN-15-2900-T.

Boussemart, Lise, Hélène Malka-Mahieu, Isabelle Girault, et al. 2014. ‘eIF4F Is a Nexus of Resistance to Anti-BRAF and Anti-MEK Cancer Therapies’. Nature 513 (7516): 105–9. 10.1038/nature13572.

Castello, Alfredo, Bernd Fischer, Katrin Eichelbaum, et al. 2012. ‘Insights into RNA Biology from an Atlas of Mammalian mRNA-Binding Proteins’. Cell 149 (6): 1393–406. 10.1016/j.cell.2012.04.031.

Chai, Peiyuan, Sina Kheiri, Andrew Kuo, et al. 2026. ‘GlycoRNA Complexed with Heparan Sulfate Regulates VEGF-A Signalling’. Nature 651 (8106): 808–18. 10.1038/s41586-025-10052-8.

Che, Yonglu, Zurab Siprashvili, Joanna R. Kovalski, et al. 2019. ‘KRAS Regulation by Small Non-Coding RNAs and SNARE Proteins’. Nature Communications 10 (1): 5118. 10.1038/s41467-019-13106-4.

Du, William W., Weining Yang, Elizabeth Liu, Zhenguo Yang, Preet Dhaliwal, and Burton B. Yang. 2016. ‘Foxo3 Circular RNA Retards Cell Cycle Progression via Forming Ternary Complexes with P21 and CDK2’. Nucleic Acids Research 44 (6): 2846–58. 10.1093/nar/gkw027.

Hentze, Matthias W., Pia Sommerkamp, Venkatraman Ravi, and Fátima Gebauer. 2025. ‘Rethinking RNA-Binding Proteins: Riboregulation Challenges Prevailing Views’. Cell 188 (18): 4811–27. 10.1016/j.cell.2025.06.021.

Lee, Sungyul, Florian Kopp, Tsung-Cheng Chang, et al. 2016. ‘Noncoding RNA NORAD Regulates Genomic Stability by Sequestering PUMILIO Proteins’. Cell 164 (0): 69–80. 10.1016/j.cell.2015.12.017.

Liau, Nicholas P. D., Avinashnarayan Venkatanarayan, John G. Quinn, et al. 2020. ‘Dimerization Induced by C-Terminal 14–3–3 Binding Is Sufficient for BRAF Kinase Activation’. Biochemistry 59 (41): 3982–92. 10.1021/acs.biochem.0c00517.

Markaki, Yolanda, Johnny Gan Chong, Yuying Wang, et al. 2021. ‘Xist Nucleates Local Protein Gradients to Propagate Silencing across the X Chromosome’. Cell 184 (25): 6174–6192.e32. 10.1016/j.cell.2021.10.022.

Mestre-Farràs, Neus, Santiago Guerrero, Nadine Bley, et al. 2022. ‘Melanoma RBPome Identification Reveals PDIA6 as an Unconventional RNA-Binding Protein Involved in Metastasis’. Nucleic Acids Research 50 (14): 8207–25. 10.1093/nar/gkac605.

Park, Eunyoung, Shaun Rawson, Kunhua Li, et al. 2019. ‘Architecture of Autoinhibited and Active BRAF–MEK1–14-3-3 Complexes’. Nature 575 (7783): 545–50. 10.1038/s41586-019-1660-y.

Peabody, D. S. 1993. ‘The RNA Binding Site of Bacteriophage MS2 Coat Protein.’ The EMBO Journal 12 (2): 595–600. 10.1002/j.1460-2075.1993.tb05691.x.

Pettersen, Eric F., Thomas D. Goddard, Conrad C. Huang, et al. 2021. ‘UCSF ChimeraX: Structure Visualization for Researchers, Educators, and Developers’. Protein Science: A Publication of the Protein Society 30 (1): 70–82. 10.1002/pro.3943.

Queiroz, Rayner M. L., Tom Smith, Eneko Villanueva, et al. 2019. ‘Comprehensive Identification of RNA-Protein Interactions in Any Organism Using Orthogonal Organic Phase Separation (OOPS)’. Nature Biotechnology 37 (2): 169–78. 10.1038/s41587-018-0001-2.

Sanchez-Vega, Francisco, Marco Mina, Joshua Armenia, et al. 2018. ‘Oncogenic Signaling Pathways in The Cancer Genome Atlas’. Cell 173 (2): 321–337.e10. 10.1016/j.cell.2018.03.035.

Sinkala, Musalula, Panji Nkhoma, Nicola Mulder, and Darren Patrick Martin. 2021. ‘Integrated Molecular Characterisation of the MAPK Pathways in Human Cancers Reveals Pharmacologically Vulnerable Mutations and Gene Dependencies’. Communications Biology 4 (1): 9. 10.1038/s42003-020-01552-6.

Siprashvili, Zurab, Dan E. Webster, Danielle Johnston, et al. 2016. ‘The Noncoding RNAs SNORD50A and SNORD50B Bind K-Ras and Are Recurrently Deleted in Human Cancer’. Nature Genetics 48 (1): 53–58. 10.1038/ng.3452.

Ule, Jernej, Kirk B. Jensen, Matteo Ruggiu, Aldo Mele, Aljaz Ule, and Robert B. Darnell. 2003. ‘CLIP Identifies Nova-Regulated RNA Networks in the Brain’. *Science (New York*, N.Y.) 302 (5648): 1212–15. 10.1126/science.1090095.

Ule, Jernej, Kirk Jensen, Aldo Mele, and Robert B. Darnell. 2005. ‘CLIP: A Method for Identifying Protein–RNA Interaction Sites in Living Cells’. *Methods*, Post-transcriptional Regulation of Gene Expression, vol. 37 (4): 376–86. 10.1016/j.ymeth.2005.07.018.

Zhu, Youming, Lei Jin, Ronghua Shi, et al. 2022. ‘The Long Noncoding RNA glycoLINC Assembles a Lower Glycolytic Metabolon to Promote Glycolysis’. Molecular Cell 82 (3): 542–554.e6. 10.1016/j.molcel.2021.11.017.

